# Transcriptomic Landscape of Hyperthyroidism in Mice Overexpressing Thyroid Stimulating Hormone

**DOI:** 10.1101/2023.10.27.564354

**Authors:** Ichiro Yamauchi, Taku Sugawa, Takuro Hakata, Akira Yoshizawa, Tomoko Kita, Yo Kishimoto, Sadahito Kimura, Daisuke Kosugi, Haruka Fujita, Kentaro Okamoto, Yohei Ueda, Toshihito Fujii, Daisuke Taura, Yoriko Sakane, Akihiro Yasoda, Nobuya Inagaki

**Affiliations:** Department of Diabetes, Endocrinology and Nutrition, Graduate School of Medicine, Kyoto University; Department of Otolaryngology-Head and Neck Surgery, Graduate School of Medicine, Kyoto University; Sugawa Clinic; Clinical Research Center, National Hospital Organization Kyoto Medical Center; Medical Research Institute KITANO HOSPITAL, PIIF Tazuke-kofukai

**Keywords:** Hyperthyroidism, Thyroid stimulating hormone, Hydrodynamic gene delivery, Thiamazole, SLC26A4

## Abstract

Hyperthyroidism is a condition with excessive thyroid hormone secretion. Activation of thyroid stimulating hormone receptor (TSHR) fundamentally leads to hyperthyroidism. The details of TSHR signaling remain to be elucidated. We conducted transcriptome analyses for hyperthyroid mice that we generated by overexpressing TSH. TSH overexpression via hydrodynamic gene delivery with pLIVE-*TSHB* and pLIVE-*CGA* vectors consistently caused hyperthyroidism and goiters for at least 4 weeks in C57BL/6J mice. RNA sequencing analysis of their thyroid glands revealed that thiamazole slightly changed the thyroid transcriptome, which reinforces a conventional theory that thiamazole decreases thyroid hormone secretion via inhibition of thyroid peroxidase activity. Meanwhile, TSH overexpression drastically changed the thyroid transcriptome. In particular, enrichment analyses identified the cell cycle, phosphatidylinositol-3 kinase/Akt pathway, and Ras-related protein 1 pathway as possibly associated with goiter development. Regarding the role of TSHR signaling in hyperthyroidism, it is noteworthy that *Slc26a4* was exclusively upregulated among genes crucial to thyroid hormone secretion at both 1 and 4 weeks after hydrodynamic gene delivery. To verify the relationship between this upregulation and hyperthyroidism, we overexpressed TSH in *Slc26a4* knockout mice. TSH overexpression caused hyperthyroidism in *Slc26a4* knockout mice, equivalent to that in control mice. To summarize, we analyzed hyperthyroid mice generated by TSH overexpression. We did not observe significant changes in known genes and pathways involved in thyroid hormone secretion. Thus, our datasets might include candidate genes that have not yet been identified as regulators of thyroid function. Our transcriptome datasets regarding hyperthyroidism can contribute to future research on TSHR signaling.

## Introduction

Hyperthyroidism is a condition in which excessive secretion of thyroid hormones elevates their circulating levels, leading to various symptoms and metabolic abnormalities. Graves’ disease is the most common cause of hyperthyroidism; its incidence is high as 20–40 cases per 100,000 population (1). First-line therapy of Graves’ disease has been conservative in the past several decades: anti-thyroid drugs (ATDs) that inhibit thyroid hormone secretion and radioactive iodine therapy. Physicians in Europe, Latin America, and Japan prefer ATDs (2). Increased use of ATDs and decreased use of radioactive iodine therapy were recently observed in the United States (3).

ATDs are used for many patients, whereas they have various adverse effects. Among major ATDs, thiamazole (MMI) and propylthiouracil (PTU) often cause liver injury and rash. They can cause agranulocytosis that progresses into life-threating infections. Furthermore, MMI should be avoided during the first trimester of pregnancy due to teratogenicity (4, 5). PTU use is not recommended for children as first-line therapy due to the high incidence of severe liver injury and vasculitis (6). Treatment of hyperthyroidism can sometimes be challenging because of these problems and insufficient effects with conventional ATDs. However, there are few promising candidates of new ATDs that inhibit thyroid hormone secretion.

The pathophysiology of hyperthyroidism can provide conceptions of new ATDs. Hyperthyroidism fundamentally develops as a result of thyroid stimulating hormone receptor (TSHR) stimulation in the thyroid gland. Autoantibodies stimulate TSHRs in Graves’ disease (1). Toxic adenoma, a thyroid neoplasm with autonomous secretion of thyroid hormones, is another cause of hyperthyroidism. It is often associated with somatic mutations in the *TSHR* gene that result in its constitutive activation (7–10). TSHR is a G protein-coupled receptor that primarily couples to G proteins that activate the protein kinase A (PKA) pathway through increasing cyclic adenosine monophosphate (cAMP) production (11). In addition, TSHR can couple to Gq/11 and activate the phospholipase C cascade with high concentration of TSH ligands (12, 13). As a result of such downstream pathways, TSHR activation increases thyroid hormone secretion and enlarges the thyroid gland, as usually seen in patients with Graves’ disease. Although the outlines of TSHR signaling are known, detailed mechanisms at the molecular level remain to be elucidated. Previous studies have identified thyroglobulin (TG), thyroid peroxidase (TPO), sodium iodide symporter (SLC5A5), type 2 iodothyronine deiodinase, and TSHR as molecules affected by TSHR activation (14–17). These were generally verified via experiments involving *in vitro* TSH administration, whereas thyroid hormone synthesis progresses in thyroid follicles consisting of follicular epithelial cells *in vivo*. It is likely that iodine concentrations in thyroid follicles are varied in hyperthyroidism because TSH increases thyroidal levels of SLC5A5, the main transporter of iodine into the thyroid gland (18). Iodine is a regulator of thyroid hormone secretion itself, which is known as the Wolff-Chaikoff effect; iodine administration represses thyroid hormone secretion with attenuation of thyroidal SLC5A5 expression (19). In this context, we believe that *in vivo* experiments to analyze cells maintaining follicular structure are necessary.

In the present study, we conducted *in vivo* experiments to understand the molecular landscape of background TSHR signaling in hyperthyroidism. Previous studies of *in vitro* TSH administration had limitations because they did not consider changes in condition of thyroid follicles. To date, *in vivo* models of hyperthyroidism have been generated by immunization of TSHR protein to replicate Graves’ disease (20), but the immunized animals do not consistently develop hyperthyroidism, even in recent models (21). Furthermore, the strategy of immunization might cause bias due to lymphocyte infiltration of the thyroid gland. We believe that hyperthyroid animals without immunization are more suitable for studying hyperthyroidism. Here, we demonstrate the phenotypes of our hyperthyroid mice generated by TSH overexpression using *in vivo* transfection and transcriptome datasets of their thyroid glands.

## Results

### Generation of hyperthyroid mice

We generated mice with hyperthyroidism through continuous activation of TSHR achieved by providing excessive concentrations of its ligand, TSH. To accomplish this strategy, we developed a method for overexpressing TSH via *in vivo* transfection with hydrodynamic gene delivery (22). TSH forms a heterodimer consisting of a specific β subunit encoded by *TSHB* (23) and an α subunit coded by *CGA* that is common to luteinizing hormone, follicle-stimulating hormone, and chorionic gonadotropin (24). Therefore, we cloned human *TSHB* and *CGA* genes into pLIVE vectors and hydrodynamically injected 25 μg of these vectors into C57BL/6J mice according to our previous studies (25–27). At 1 week after hydrodynamic gene delivery, human TSH was not detected in the serum when each vector was separately injected, but TSH was strongly detected when both vectors were co-injected (Supplementary Figure 1A). Furthermore, we observed elevated concentrations of serum free thyroxine (fT4) and free triiodothyronine (fT3) in co-injected mice (Supplementary Figure 1B, 1C).

Thereafter, the effects of TSH overexpression based on transfection with both pLIVE-*TSHB* and pLIVE-*CGA* vectors were further verified at 1 week after hydrodynamic gene delivery. We determined the dose-dependent effects by analyzing 3 groups based on the amount of injected vectors (Figure 1A). Human TSH was detected in the serum of the 5 μg group and at higher levels in the 25 μg group (Figure 1B). Dose-dependent increases were also observed in serum levels of fT4 and fT3 (Figure 1C, 1D), which was further verified by their positive correlations with serum human TSH levels: fT4 and TSH, r = 0.74 (*p* < 0.001) and fT3 and TSH, r = 0.77 (*p* < 0.001) (Supplementary Figure 1D, 1E).

**Figure 1.**
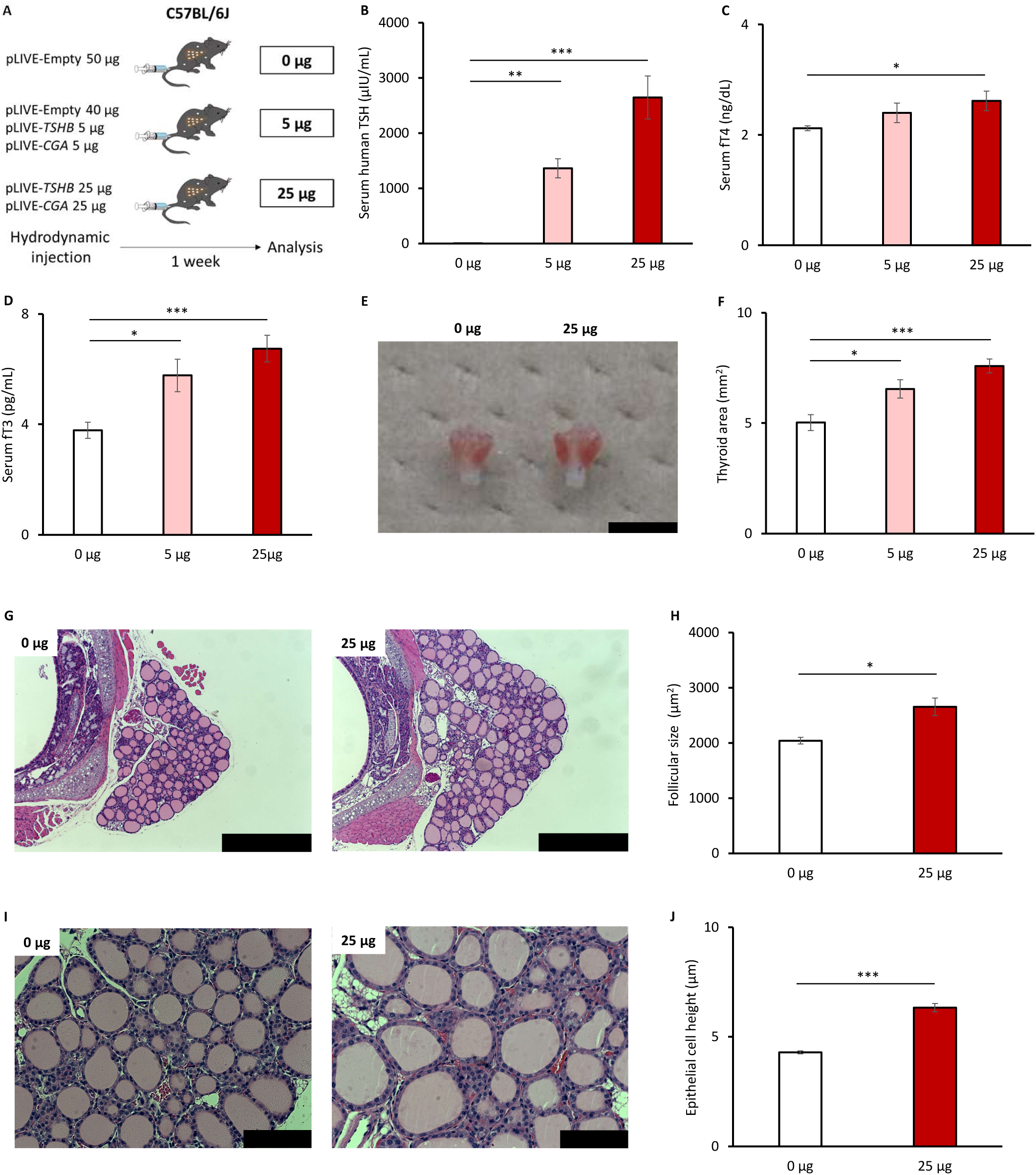
Phenotypes of mice overexpressing thyroid stimulating hormone (TSH) at 1 week after hydrodynamic gene delivery. (A) Schema for the experimental design by dose of vectors injected. *n* = 8 for each group. (B–D) Serum levels of thyroid hormones. fT4, free thyroxine; fT3, free triiodothyronine. (E) Gross appearance of thyroid glands. The black scale bar is 5 mm. (F) Thyroid gland size measured as thyroid area based on photographs. (G) Histological images of thyroid glands with hematoxylin and eosin staining at 50X magnification. The black scale bar is 500 μm. (H) Mean follicle size calculated based on histological images. (I) Histological images of thyroid glands with hematoxylin and eosin staining at 200X magnification. The black scale bar is 100 μm. (J) Epithelial cell height measured based on histological images. Data are represented as means ± standard error of the mean (SEM). Statistical analyses were performed using one-way analysis of variance (ANOVA) followed by the Dunnett test with comparison to the 0 μg group for panels B–D and F and using Student’s *t*-test for panels H and J. \**p* < 0.05, \*\**p* < 0.01, and \*\*\**p* < 0.001.

In addition to these expected changes in the serum, we found enlargement of the thyroid gland after transfection (Figure 1E). Goiters developed in a dose-dependent manner (Figure 1F); goiter severity was positively correlated with serum human TSH levels (r = 0.53, *p* < 0.05) (Supplementary Figure 1F). Histological changes in the thyroid gland included increases in follicle size (Figure 1G, 1H) and follicular epithelial cell height (Figure 1I, 1J).

Subsequently, we evaluated the persistence of induced hyperthyroidism by analyzing mice at 4 weeks after hydrodynamic gene delivery (Figure 2A). Human TSH was still detected in a dose-dependent manner (Figure 2B). Accordingly, increases in serum levels of fT4 and fT3 persisted (Figure 2C, 2D). Goiter development was also seen in a dose-dependent manner (Figure 2E, 2F). Serum human TSH levels were positively correlated with serum levels of fT4 and fT3 (fT4 and TSH, r = 0.81 (*p* < 0.001); fT3 and TSH, r = 0.65 (*p* < 0.001) and with goiter severity (thyroid area and TSH, r = 0.78 (*p* < 0.001)) (Supplementary Figure 1G–1I). Histological changes after 4 weeks were partially inconsistent with those after 1 week. Follicle size was similarly increased (Figure 2G, 2H), but follicular epithelial cell height was decreased (Figure 2I, 2J).

**Figure 2.**
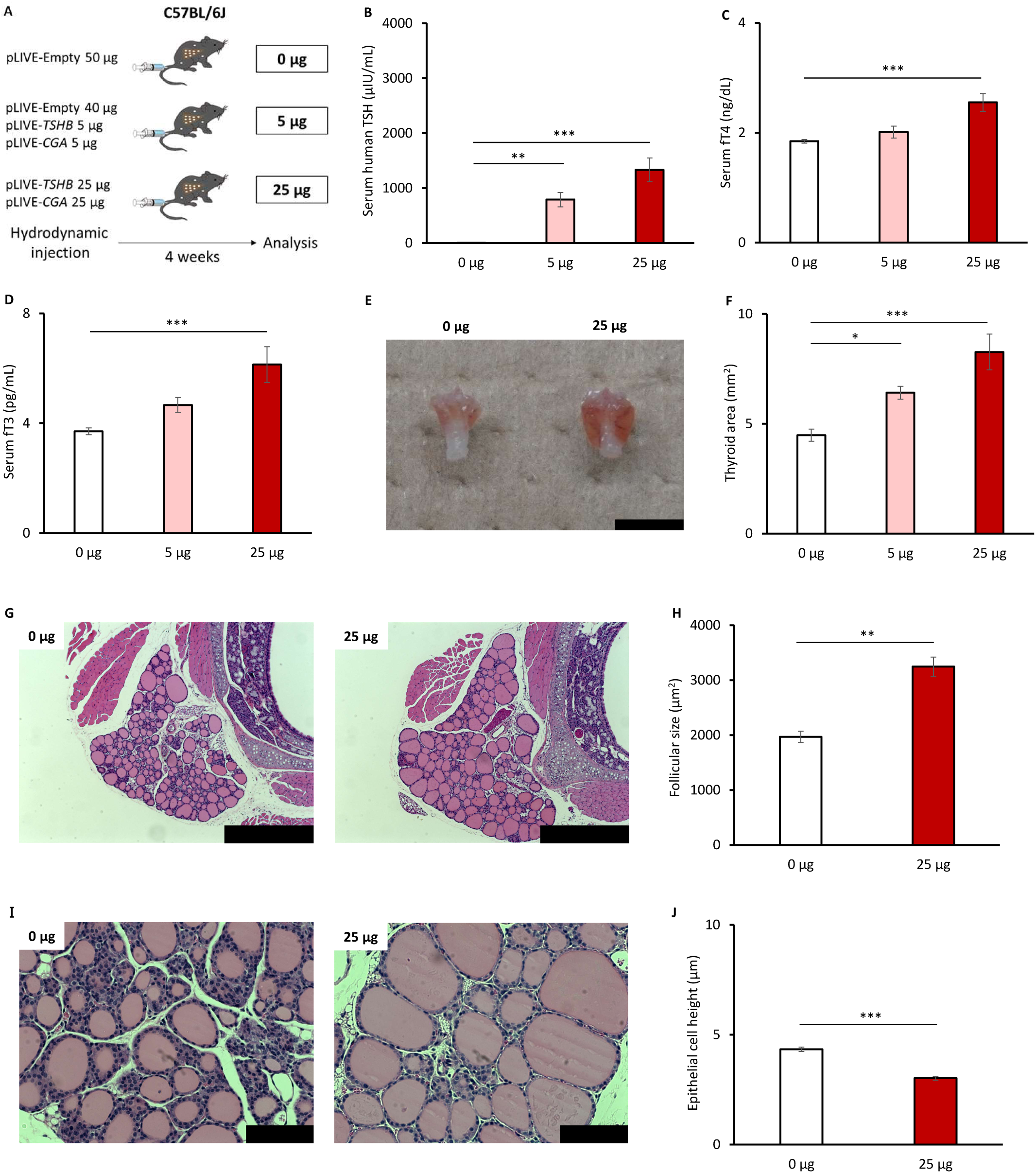
Phenotypes of mice overexpressing TSH at 4 weeks after hydrodynamic gene delivery. (A) Schema for the experimental design by dose of vectors injected. *n* = 8 for each group. (B–D) Serum levels of thyroid hormones. (E) Gross appearance of thyroid glands. The black scale bar is 5 mm. (F) Thyroid gland size measured as thyroid area based on photographs. (G) Histological images of thyroid glands with hematoxylin and eosin staining at 50X magnification. The black scale bar is 500 μm. (H) Mean follicle size calculated based on histological images. (I) Histological images of thyroid glands with hematoxylin and eosin staining at 200X magnification. The black scale bar is 100 μm. (J) Epithelial cell height measured on histological images. Data are represented as means ± SEM. Statistical analyses were performed using ANOVA followed by the Dunnett test with comparison to the 0 μg group for panels B–D and F and using Student’s *t*-test for panels H and J. \**p* < 0.05, \*\**p* < 0.01, and \*\*\**p* < 0.001.

### Thyroid transcriptome of hyperthyroid mice

Our hyperthyroid mice exhibited excessive secretion of thyroid hormones and goiter development through the effects of TSH overexpression. Using this model, we subsequently obtained data on the thyroid transcriptome using RNA sequencing (RNA-seq). In addition to investigating TSHR signaling, we aimed to clarify mechanisms of ATD action to obtain therapeutic targets for hyperthyroidism. We designed the experiment as follows (Figure 3A): the TSH group was injected with 25 μg each of the pLIVE-*TSHB* and pLIVE-*CGA* vectors and the Empty group was injected with 50 μg of the pLIVE-Empty vector. The mice were further treated with MMI in their drinking water and analyzed after 1 week.

**Figure 3.**
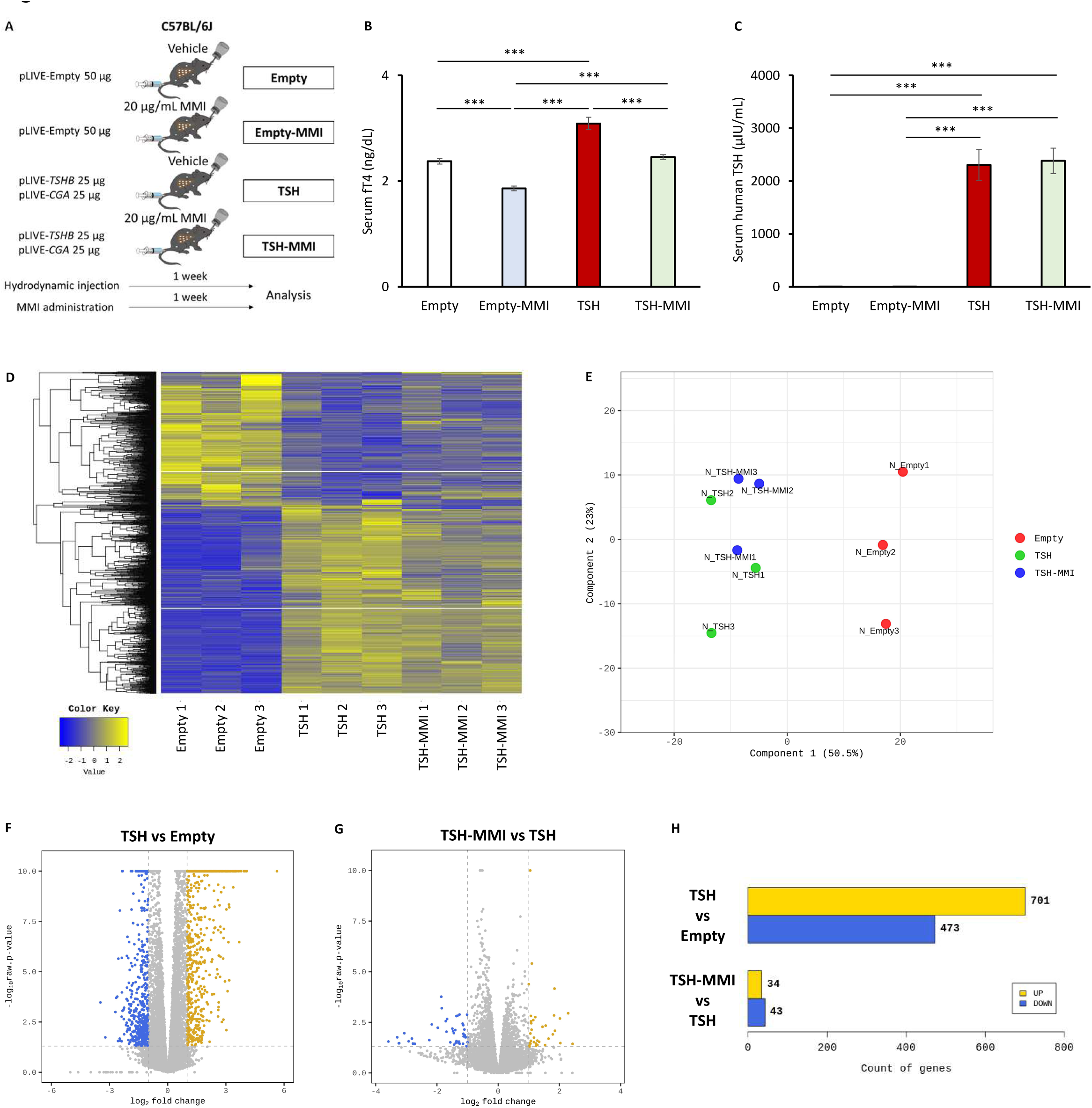
Transcriptome analyses of the thyroid glands of mice overexpressing TSH at 1 week after hydrodynamic gene delivery. (A) Schema for the experimental design. Thiamazole (MMI) was continuously administered via drinking water at 20 μg/mL. (B, C) Serum levels of thyroid hormones. *n* = 6 each. (D–H) Summary of results of RNA sequencing (RNA-seq). *n* = 3 each, selected based on proximity of serum fT4 levels to the average of each group. (D) Hierarchical clustering, (E) principal component analysis, (F) volcano plot of differentially expressed genes (DEGs) comparing the TSH and Empty groups, (G) volcano plot of DEGs comparing the TSH-MMI and TSH groups, and (H) numbers of DEGs in each comparison. Data are represented as means ± SEM. Statistical analyses were performed using ANOVA followed by the Tukey-Kramer test. \*\*\**p* < 0.001.

Hyperthyroidism was reproducibly achieved in the TSH group, which was seen as increases in serum levels of fT4 and fT3 (Figure 3B, Supplementary Figure 2A). We could match serum fT4 levels of the TSH-MMI group with those of the Empty group to avoid biases arising from differences in circulating levels of thyroid hormones (Figure 3B). TSH was equivalently detected in the serum of the TSH and TSH-MMI groups (Figure 3C).

The RNA-seq dataset of their thyroid glands is presented in Supplementary Table 1. Clustering and principal component analyses revealed clear differences between the Empty and TSH groups, whereas slight differences were observed between the TSH and TSH-MMI groups (Figure 3D, 3E). As for differentially expressed genes (DEGs), TSH overexpression substantially upregulated 701 genes and downregulated 473 genes, while MMI administration in presence of TSH overexpression only upregulated 34 genes and downregulated 43 genes (Figure 3F–3H).

In addition, similar analyses were performed at 4 weeks after hydrodynamic gene delivery (Figure 4A). MMI was only administered during the last week. Induction of hyperthyroidism was verified; the TSH group had increased serum levels of fT4 and fT3 (Figure 4B, Supplementary Figure 2B). Similarly to the result at 1 week, MMI administration decreased serum fT4 levels in the TSH-MMI group equivalent to those of the Empty group (Figure 4B). TSH overexpression was confirmed based on elevated serum human TSH levels (Figure 4C).

**Figure 4.**
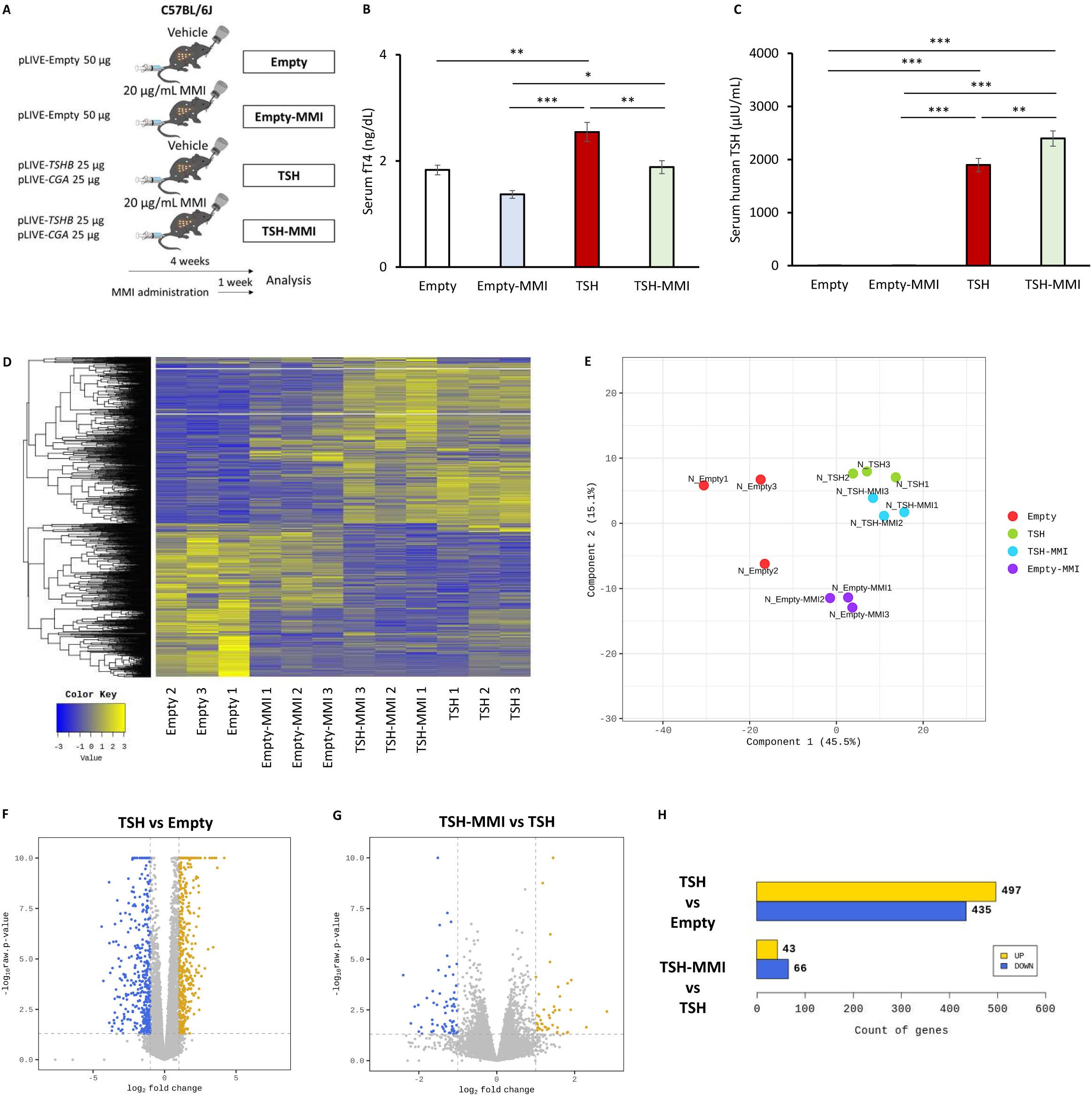
Transcriptome analyses of the thyroid glands of mice overexpressing TSH at 4 weeks after hydrodynamic gene delivery. (A) Schema for the experimental design. MMI was administered only for the last week. (B, C) Serum levels of thyroid hormones. *n* = 6 each. (D–H) Summary of results of RNA-seq. *n* = 3 each, selected based on proximity of serum fT4 levels to the average of each group. (D) Hierarchical clustering, (E) principal component analysis, (F) volcano plot of DEGs comparing the TSH and Empty groups, (G) volcano plot of DEGs comparing the TSH-MMI and TSH groups, and (H) numbers of DEGs in each comparison. Data are represented as means ± SEM. Statistical analyses were performed using ANOVA followed by the Tukey-Kramer test. \**p* < 0.05, \*\**p* < 0.01, and \*\*\**p* < 0.001.

At 4 weeks, we performed RNA-seq for all 4 groups (Supplementary Table 2). Clustering and principal component analyses revealed differences between the Empty and TSH groups as well as between the Empty and Empty-MMI groups, whereas no clear differences were seen between the TSH and TSH-MMI groups (Figure 4D, 4E). In terms of DEGs, TSH overexpression substantially upregulated 497 genes and downregulated 435 genes, while MMI administration in the presence of overexpressed TSH only upregulated 43 genes and downregulated 66 genes (Figure 4F–4H). On the other hand, MMI administration without TSH overexpression upregulated 143 genes and downregulated 236 genes (Supplementary Figure 2C, 2D). We picked 17 genes crucial to thyroid hormone secretion, referred to as thyroidal genes, and examined their changes. MMI administration in presence and absence of TSH overexpression altered few thyroidal genes (Supplementary Figure 2E, 2F). As we have found small changes in the thyroid transcriptome with MMI administration, we focused on comparisons between the TSH and Empty groups in following analyses.

Enrichment analyses were performed using gene ontology (GO) and the Kyoto Encyclopedia of Genes and Genomes (KEGG). We compared the TSH group with the Empty group. Results at 1 week are shown in Supplementary Figure 3 and those at 4 weeks are in Supplementary Figure 4. At 1 week, terms related to the cell cycle were often seen with TSH overexpression (Supplementary Figure 3). Although terms related to muscle were often seen with TSH overexpression at 4 weeks (Supplementary Figure 4), they were also seen with MMI administration without TSH overexpression at 4 weeks (Supplementary Figure 5), which was difficult to comprehend. Through these analyses, we could not find remarkable terms related to hyperthyroidism.

We examined individual DEGs and listed the top 30 genes upregulated with TSH overexpression. At 1 week, we found upregulated genes associated with the cell cycle such as *Cdkn3*, *Plk1*, and *Ccnb1*, which was consistent with the results of enrichment analyses (Table 1). It was noteworthy that *Slc26a4* had the second highest change at 1 week (Table 1). Upregulation of *Slc26a4* was similarly observed at 4 weeks (Table 2). We reviewed the datasets for thyroidal genes, which revealed that *Slc26a4* was exclusively upregulated at both 1 and 4 weeks (Figure 5).

**Figure 5.**
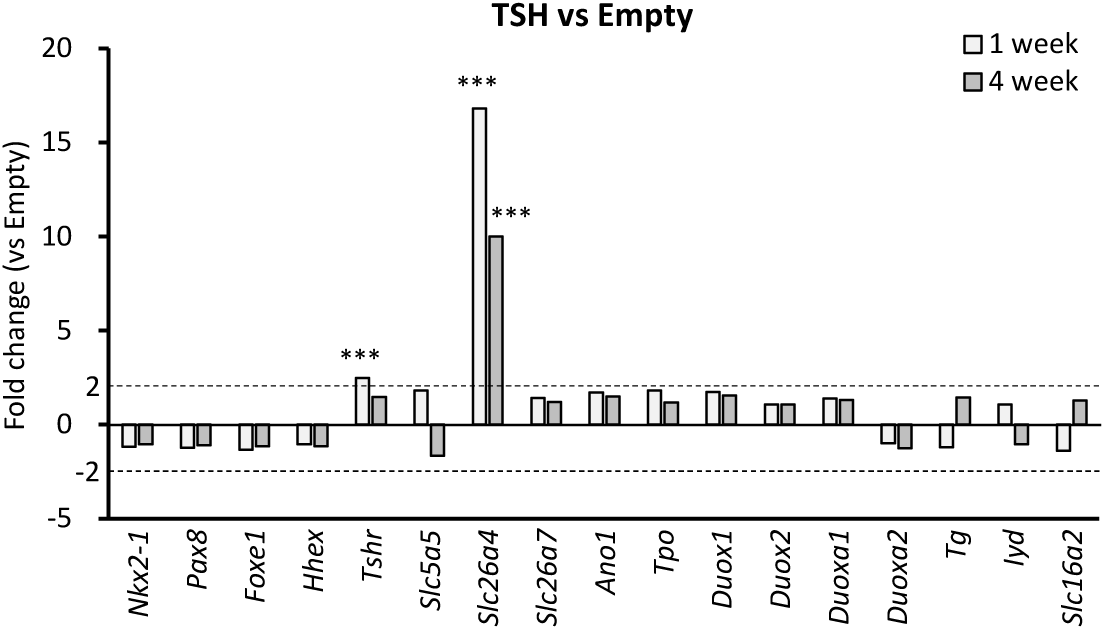
Summary of RNA-seq regarding thyroidal genes. See Supplementary Tables 1 and 2 for the source data. Data are represented as fold changes between the TSH group and the Empty group in transcripts per kilobase million (TPM) of genes. The Wald test was used to compare TPM of genes from each cohort. \*\*\**p* < 0.001.

**Table 1.**
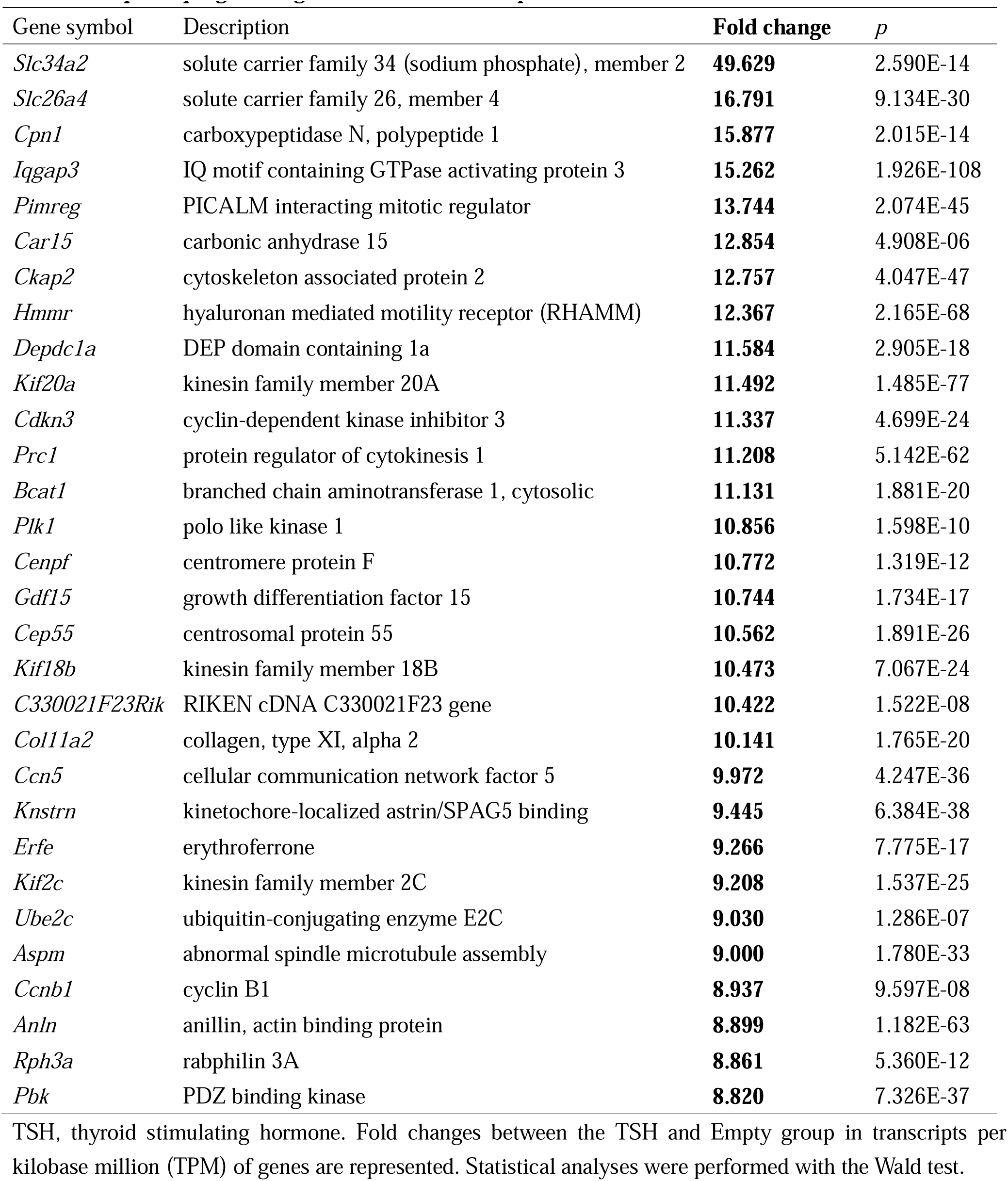
Top 30 upregulated genes with TSH overexpression at 1 week.

**Table 2.**
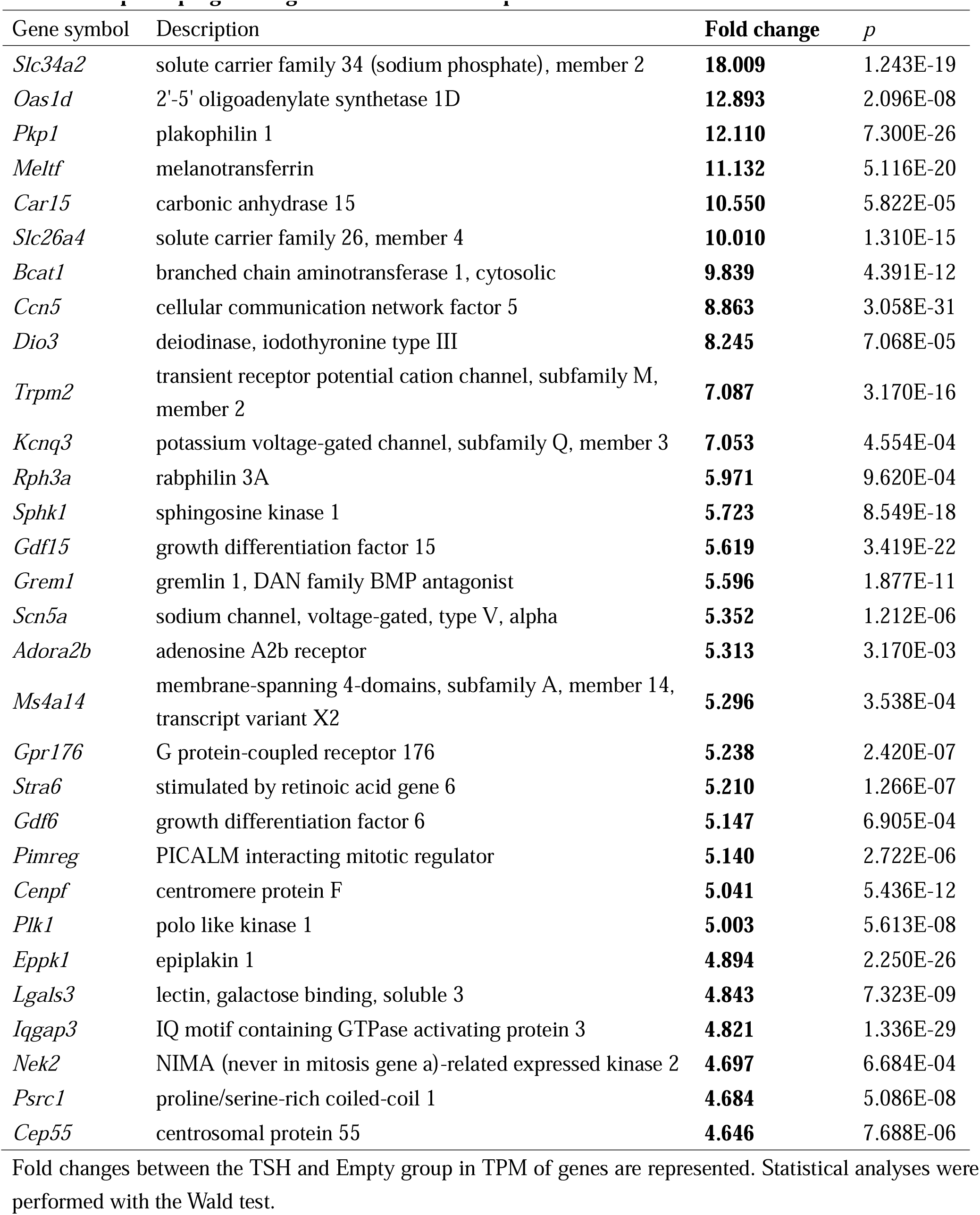
Top 30 upregulated genes with TSH overexpression at 4 weeks.

### TSH overexpression in Slc26a4 knockout mice

*Slc26a4* codes the SLC26A4 protein, also known as pendrin. SLC26A4 is an apical iodine transporter that transports iodine into thyroid follicles, which is important for thyroid hormone secretion. Attenuation of SLC26A4 action can cause hypothyroidism. Loss-of-function mutations in *SLC26A4* are known to cause Pendred syndrome, which includes hypothyroidism (28). We examined the effects of increased SLC26A4 expression on hyperthyroidism. Our working hypothesis was that hyperthyroidism is attenuated if SLC26A4 expression is not upregulated in response to TSH overexpression.

To test this hypothesis, we analyzed *Slc26a4* knockout mice. More specifically, we caused hyperthyroidism via TSH overexpression in heterozygous (Hetero-TSH group) and homozygous *Slc26a4* knockout mice (Homo-TSH group). We compared them with mice injected with pLIVE-Empty vector (Hetero-Empty and Homo-Empty groups) (Figure 6A). The concentration of vectors injected was doubled because *Slc26a4* knockout mice somehow exhibited relatively low serum levels of human TSH with TSH overexpression (Figure 6B). Nevertheless, the Hetero-TSH and Homo-TSH groups certainly developed hyperthyroidism and goiters (Figure 6C–6F). No significant differences in serum levels of fT4, fT3, or human TSH were observed between the Hetero-TSH and Homo-TSH groups (Figure 6B–6D). Goiter severity was also similar between the Hetero-TSH and Homo-TSH groups (Figure 6E, 6F). Although follicle size and follicular epithelial cell height were significantly increased with TSH overexpression, no differences were observed between the Hetero-TSH and Homo-TSH groups (Figure 6G–6J).

**Figure 6.**
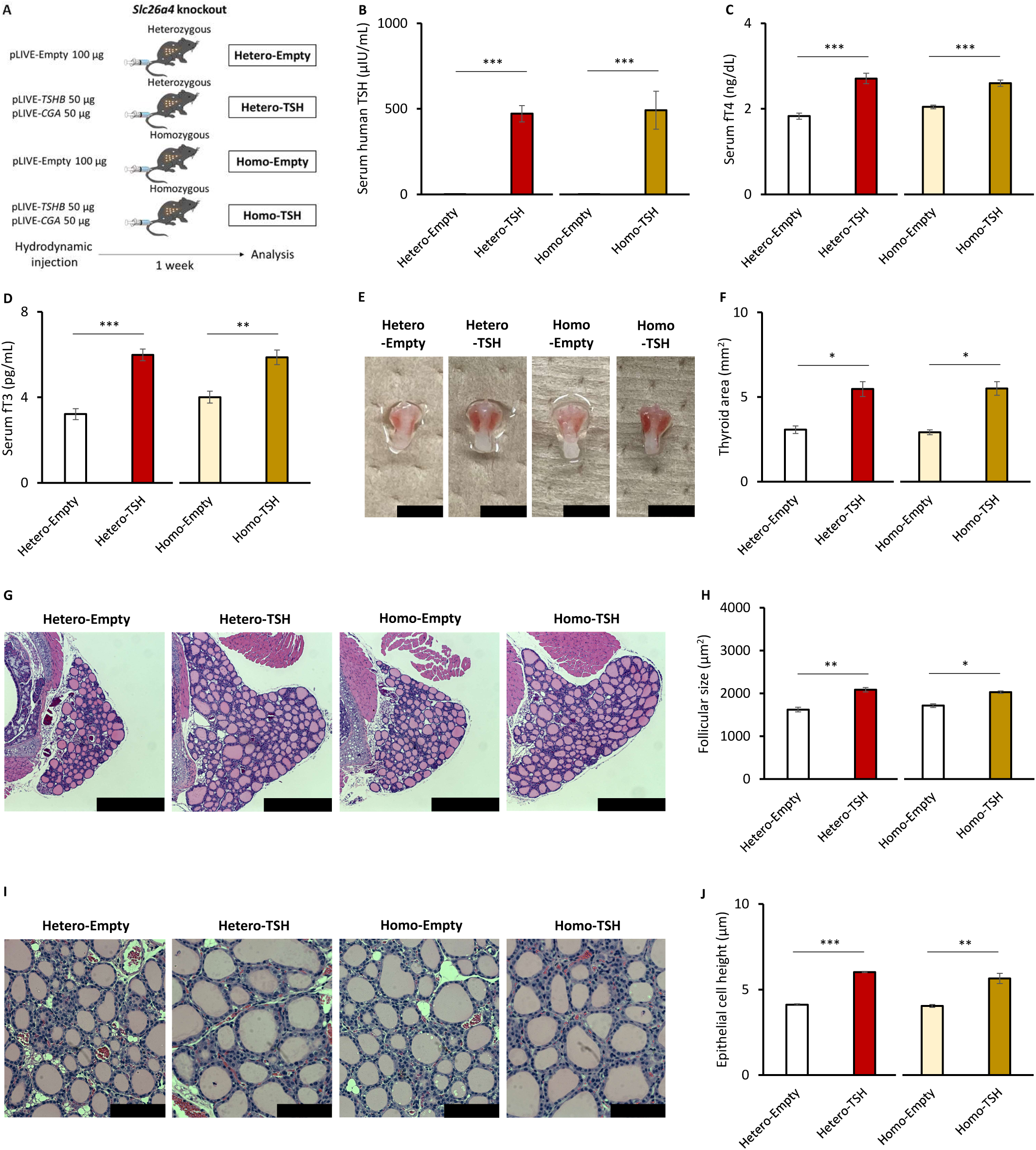
Phenotypes of *Slc26a4* knockout mice overexpressing TSH. (A) Schema for the experimental design. (B–D) Serum levels of thyroid hormones. Hetero-Empty, Hetero-TSH, *n* = 12 each; Homo-Empty, Homo-TSH, *n* = 8 each. (E) Gross appearance of thyroid glands. The black scale bar is 5 mm. (F) Thyroid gland size measured based on photographs. *n* = 3 each. (G) Histological images of thyroid glands with hematoxylin and eosin staining at 50X magnification. The black scale bar is 500 μm. (H) Mean follicle size calculated based on histological images. *n* = 3 each. (I) Histological images of thyroid glands with hematoxylin and eosin staining at 200X magnification. The black scale bar is 100 μm. (J) Epithelial cell height measured based on histological images. *N* = 3 each. Data are represented as means ± SEM. Statistical analyses were performed using Student’s *t*-test to compare the Empty and TSH groups for each knockout mouse. \**p* < 0.05, \*\**p* < 0.01, and \*\*\**p* < 0.001.

We carefully verified these results through the following examinations. In heterozygous *Slc26a4* knockout mice, there was a significant increase in *Slc26a4* mRNA levels in the thyroid gland with TSH overexpression (Supplementary Figure 6A, 6B). Examination of selected thyroidal genes revealed lower mRNA levels of *Tg*, *Tpo*, and *Slc5a5* with TSH overexpression in heterozygous *Slc26a4* knockout mice, while no significant changes were observed in homozygous *Slc26a4* knockout mice (Supplementary Figure 6C, 6D). This result raised a concern that downregulation of *Tg*, *Tpo*, and *Slc5a5* attenuates hyperthyroidism induction in heterozygous *Slc26a4* knockout mice, regardless of SLC26A4 levels.

To address this concern and confirm reproducibility, we performed additional analyses at 3 days after hydrodynamic gene delivery (Supplementary Figure 7A). Both heterozygous and homozygous *Slc26a4* knockout mice had increases in serum human TSH levels with TSH overexpression (Supplementary Figure 7B) and developed equivalent degrees of hyperthyroidism as determined based on serum fT4 levels (Supplementary Figure 7C). Serum fT3 levels were also significantly increased in heterozygous *Slc26a4* knockout mice and insignificantly increased in homozygous *Slc26a4* knockout mice (Supplementary Figure 7D). Increases in *Slc26a4* mRNA levels with TSH overexpression in heterozygous *Slc26a4* knockout mice were also confirmed (Supplementary Figure 7E, 7F). Homozygous *Slc26a4* knockout mice did not have *Slc26a4* mRNA (Supplementary Figure 7G). However, SLC26A4 protein was difficult to detect in the thyroid glands of heterozygous *Slc26a4* knockout mice (Supplementary Figure 7H). We could not detect SLC26A4 protein in the thyroid glands of heterozygous *Slc26a4* knockout mice with TSH overexpression, even though *Slc26a4* mRNA levels were increased (Supplementary Figure 7I). These results suggested that SLC26A4 protein is slightly or not expressed in the murine thyroid gland. Examination of selected thyroidal genes revealed decreases in levels of *Tg* and *Slc5a5* mRNA with TSH overexpression in both heterozygous and homozygous *Slc26a4* knockout mice, while other genes were not changed (Supplementary Figure 8A, 8B).

## Discussion

In the present study, we generated hyperthyroid mice through TSH overexpression with hydrodynamic gene delivery using both pLIVE-*TSHB* and pLIVE-*CGA* vectors. The hyperthyroid mice consistently developed hyperthyroidism and goiters for at least 4 weeks. We conducted transcriptome analysis on their thyroid glands to elucidate TSHR signaling. Although few changes in the thyroid transcriptome were observed when the hyperthyroid mice were treated with MMI, drastic changes were induced with TSH overexpression at both 1 and 4 weeks after hydrodynamic gene delivery. We found that *Slc26a4* is the only upregulated thyroidal gene at both 1 and 4 weeks. To examine whether SLC26A4 is associated with the development of hyperthyroidism, we caused hyperthyroidism in *Slc26a4* knockout mice. *Slc26a4* knockout mice certainly developed hyperthyroidism through TSH overexpression, but had similar degrees of hyperthyroidism as control mice.

Several murine models of hyperthyroidism have been generated through immunization with TSHR protein (20, 21). Interestingly, Jaeschke *et al.* generated mice with a knocked-in human *TSHR* sequence that had a constitutively active mutation (29). Serum fT4 levels were higher in the homozygous *TSHR* knock-in mice. Goiters developed in both the heterozygous and homozygous *TSHR* knock-in mice. Hyperthyroidism developed at 1 month of age while goiters developed at 2 months of age. Our hyperthyroid mice and *TSHR* knock-in mice were similar in that both had TSHR activation and developed hyperthyroidism with goiters.

Our hyperthyroid mouse, generated by overexpressing human TSH, has the following merits compared to previous models. First, hyperthyroidism consistently and promptly developed at 3 days after hydrodynamic gene delivery and persisted for at least 4 weeks. Second, we can validate and monitor TSH overexpression by measuring serum human TSH levels. Compared with human TSH, murine TSH levels are difficult to measure (30). The kit we used could detect overexpression of human TSH. There was no significant cross-reaction with endogenous murine TSH, as seen in the results of the Empty group. Third, the strategy of acquired generation can avoid bias arising during growth periods. Lastly, we can cause hyperthyroidism in various mice because one hydrodynamic injection of the same plasmid vectors is just needed. We believe our technique of hyperthyroidism induction can be also useful in other investigations.

Hydrodynamic gene delivery is a technique for *in vivo* transfection. Prolonged overexpression is obtained by using a pLIVE vector backbone. In a previous study, we applied this system to increase circulating levels of C-type natriuretic peptide and verified the prolonged increase even at 4 weeks after transfection (25). Attempts to increase circulating protein levels with hydrodynamic gene delivery using pLIVE vectors were performed for fibroblast growth factor 21 (31, 32) and soluble T-cadherin (33). Kwon *et al*. demonstrated that elevation of plasma fibroblast growth factor 21 levels was maintained at 100 days after hydrodynamic injection (32). Thus, our hyperthyroid mice are also expected to maintain high serum TSH levels for several months. As the *TSHR* knock-in mice developed papillary thyroid carcinomas at 12 months of age (29), long-term observations of our hyperthyroid mice may yield interesting findings.

Transcriptome analyses of the thyroid gland focusing on thyroid dysfunction have been limited to reports on a murine model of Graves’ disease (34) and a murine model of iodine treatment (35). To the best of our knowledge, there have been no clinical reports about the thyroid transcriptome of patients with hyperthyroidism. In the present study, our hyperthyroid mice provided valuable information on the molecular signature of hyperthyroidism. First, we investigated mechanisms of MMI action on hyperthyroidism but could find few changes in the thyroid transcriptome. In other words, MMI seemed to attenuate hyperthyroidism regardless of transcriptional regulation. ATDs such as MMI and PTU inhibit TPO activity (36, 37). Our results reinforce a conventional theory that ATDs decrease thyroid hormone secretion by inhibiting TPO (38).

GO enrichment analyses of TSH overexpression at 1 week shed light on the cell cycle (Supplementary Figure 3). *Cdkn3*, *Plk1*, and *Ccnb1,* which are related to the cell cycle, were listed among the top 30 upregulated genes at 1 week (Table 1). It is well known that TSH activates cell proliferation (11), which might be involved in cell cycle progression (39, 40). Meanwhile, changes related to the cell cycle were not seen in GO terms or the top 30 upregulated genes at 4 weeks (Supplementary Figure 4, Table 2). Instead, we noticed the phosphatidylinositol-3 kinase (PI3K)/Akt pathway and Ras-related protein 1 (Rap1) pathway among KEGG terms (Supplementary Figure 3, Supplementary Figure 4). Previous studies have verified that TSHR signaling includes activation of PI3K and consequent phosphorylation of Akt (41–43). TSHR signaling directly activates the phospholipase C cascade as well as cAMP/PKA pathway via G proteins (12, 13), but how these mechanisms contribute to PI3K/Akt pathway activation is unclear. PI3K inhibition attenuates the proliferative effects of TSH (43, 44), which support involvement of the PI3K/Akt pathway in goiter development in our hyperthyroid mice.

On the other hand, Rap1 induction by TSH through both PKA-dependent and PKA-independent pathways has been reported (45, 46). Rap1 is known as an activator of mitogen-activated protein kinase signaling through BRAF activation (47). The similar action of Rap1 was observed in the FRTL5 rat thyroid cell line (48). Given that induction of a gain-of-function BRAF mutation enlarged murine thyroid glands by promoting mitosis (49), we believe that Rap1 signaling also contributed to goiter development of our hyperthyroid mice.

We also examined whether SLC26A4 played a significant role in the development of hyperthyroidism because its encoding gene, *Slc26a4*, was exclusively upregulated among thyroidal genes with TSH overexpression. *Slc26a4* mRNA was also upregulated in a murine model of Graves’ disease (34). Pesce *et al.* reported that TSH regulates SLC26A4 abundance in the plasma membrane in the PCCL-3 rat thyroid cell line (50). In addition, examinations of human thyroid tissues have revealed that SLC26A4 expression is augmented in Graves’ disease (51, 52). Hypothyroidism due to loss-of-function mutations in *SLC26A4* occur in Pendred syndrome (28), but the effects of increased levels of SLC26A4 on thyroid function have not been clarified. In this context, we generated and analyzed *Slc26a4* knockout mice with hyperthyroidism that did not have increases in *Slc26a4* mRNA expression with TSH overexpression. SLC26A4 depletion did not affect their thyroid function and morphology. We excluded compensative changes in the gene expression of components necessary for thyroid hormone secretion including *SLC5A5*, *ANO1*, *SLC26A7*, and other iodine transporters (53–55). Furthermore, chronological adaptation to hyperthyroidism was ruled out based on earlier analyses, at 3 days after hydrodynamic gene delivery. This finding might be explained with a report that *Slc26a7* knockout mice develop hypothyroidism but *Slc26a4* knockout mice do not (56). As seen in our western blot results, SLC26A4 protein might be slightly expressed and not play a significant role in the murine thyroid gland.

SLC26A4 did not seem to be involved in hyperthyroidism, which was interesting because our mice developed hyperthyroidism without any changes in known thyroidal genes. In the process of developing hyperthyroidism through activation of TSHR signaling, a number of genes should be regulated because PKA generally phosphorylates cAMP response element binding protein and regulates its target genes (57, 58). In other words, DEGs in our datasets might include candidate genes that have not yet been identified as regulators of thyroid function.

We hope that our RNA-seq datasets (Supplementary Table 1, Supplementary Table 2) will provide inspiration to other researchers. We manually reviewed the datasets and found the following suggestions. *Lgals3*, a gene encoding galectin-3, were upregulated. Galectin-3 is a well-known marker for differentiated thyroid cancer (59). Considering that TSH suppression therapy is effective against differentiated thyroid cancer, a relationship between TSHR signaling and galectin-3 is reasonable. Moreover, *Gdf15* was seen among the top 30 genes at both 1 and 4 weeks. Growth differentiation factor 15 (GDF15) is associated with various endocrine conditions (60). Patients with hyperthyroidism have high serum GDF15 levels (61), as well as patients with papillary thyroid carcinoma (62). In addition, the papillary thyroid carcinomas often express GDF15 (62). As for hyperthyroidism, we are interested in *Siglec1* upregulation. Hashimoto *et al.* reported that patients who experience relapse of Graves’ disease have high *SIGLEC1* mRNA levels in peripheral leukocytes (63). The role of sialic acid-binding immunoglobulin-like lectin-1 (SIGLEC1) in the thyroid gland has not been clarified. Perspectives on TSHR signaling might contribute to greater understanding of thyroid biology. To provide readers valuable information on the thyroid transcriptome, Supplementary Table 3 includes a list of genes upregulated at both 1 and 4 weeks.

The present study has several limitations. First, the TSH overexpressed in our system might not have post-translational modifications because it was predominantly produced in the liver (26), not the pituitary gland, which is different from physiological TSH secretion. Likewise, TSHR-stimulating antibodies in Graves’ disease might affect TSHR signaling differently. Second, differences in thyroid physiology between species should be considered. Although thyroidal genes were not rarely changed with TSH overexpression in our mice (Figure 5), the human thyroid gland might undergo different changes. For example, TSH administration increases mRNA levels of *TG*, *TPO*, and *SLC5A5* of human thyrocytes in primary culture (17). Analyses with human thyroid tissues or organoids are needed to address this limitation. Third, we analyzed the thyroid transcriptome using bulk RNA-seq, which included C cells and the parathyroid glands as well as follicular epithelial cells due to technical difficulties. We actually found decreases in their respective marker genes, *Calca* and *Pth*, with TSH overexpression (Supplementary Figure 1, Supplementary Figure 2). We hypothesize that TSH overexpression increased the amount of RNA originating from follicular epithelial cells and caused the apparent decrease in the amount of RNA from non-follicular epithelial cells. Therefore, interpretation of downregulated genes might be difficult. Single-cell analysis or other techniques are required for further investigation.

In conclusion, we generated a mouse model of hyperthyroidism based on TSH overexpression to clarify the molecular landscape of TSHR signaling in a hyperthyroid state. Although MMI administration caused few changes in the thyroid transcriptome, TSH overexpression induced drastic changes. As *Slc26a4* was exclusively upregulated among known thyroidal genes with TSH overexpression, we verified the significance of *Slc26a4* in hyperthyroidism based on analyses of *Slc26a4* knockout mice. As a result, *Slc26a4* knockout mice developed hyperthyroidism similar to control mice. It is possible that unrecognized genes are involved in the development of hyperthyroidism. We hope the transcriptome datasets presented here will contribute to future research.

## Materials and Methods

### Plasmid construction

We cloned the coding sequences of human *TSHB* and *CGA* into pLIVE vectors (Mirus Bio, Madison, WI) using a polymerase chain reaction (PCR) technique with primers containing restriction enzyme sites and the Kozak sequence (Supplementary Table 4). The constructed vectors were verified based on sequencing. We prepared a large amount of these plasmid vectors using a NucleoBond Xtra Midi Plus kit (Macherey-Nagel, Düren, Germany). Concentrations of plasmid DNA were measured with a Nanodrop-1000 spectrophotometer (Thermo Fisher Scientific, Waltham, MA).

### Animals

We purchased male C57BL/6J mice from Japan SLC, Inc. (Hamamatsu, Japan). In the present study, *Slc26a4* knockout mice (64, 65) were backcrossed with C57BL/6J mice. All experimental procedures involving animals were approved by the Animal Research Committee of the Kyoto University Graduate School of Medicine (approval number, Med Kyo 19512). Animal care and all animal experiments were conducted in accordance with our institutional guidelines. All animals were housed at 23°C in a 14:10–hour light/dark cycle.

### Hydrodynamic gene delivery and MMI administration

Six-week-old male C57BL/6J mice were injected with 50 μg of pLIVE-Empty vector or a mixture of 25 µg of pLIVE-*TSHB* vector and 25 µg of pLIVE-*CGA* vector using a hydrodynamic gene delivery method, as in our previous studies (25–27). Briefly, a solution of plasmid DNA and sterilized saline (Otsuka Pharmaceutical Factory, Tokyo, Japan) was prepared as a total volume equivalent to 8–10% of the weight of each mouse. The solution was rapidly injected within 10 seconds into the tail vein using a 27-gauge needle (Terumo Corporation, Tokyo, Japan). Male *Slc26a4* knockout mice were injected with 100 μg of pLIVE-Empty vector or a mixture of 50 µg of pLIVE-*TSHB* vector and 50 µg of pLIVE-*CGA* vector at 6 weeks of age. Regarding administration of MMI (Tokyo Chemical Industry Co., Ltd., Tokyo Japan), mice had *ad libitum* access to drinking water containing 20 µg/mL of MMI.

### Sample collection

At 3 days, 1 week, or 4 weeks after hydrodynamic gene delivery, we sacrificed the mice and collected samples. All mice were sacrificed using isoflurane exposure at the end of the light cycle. To obtain serum, blood samples were collected in microtubes and left undisturbed for 45 minutes at room temperature. Next, these samples were left on ice and immediately centrifuged at 3,000 g for 15 minutes at 4°C. The supernatants were collected and stored at −80°C until use. Thyroid glands harvested for RNA extraction were preserved in RNAprotect Tissue Reagent (QIAGEN, Venlo, Netherlands) at −20°C until use. Thyroid glands harvested for histological analyses were first fixed in 4% paraformaldehyde phosphate buffer solution (FUJIFILM Wako Pure Chemical Corporation, Osaka, Japan), embedded in paraffin, and stained using hematoxylin and eosin. Thyroid glands harvested for western blotting analyses were collected in microtubes, immediately frozen in liquid nitrogen, and stored at −80°C until use.

### Thyroid morphological analysis

Morphological analysis of thyroid glands was performed using ImageJ software (National Institutes of Health, Bethesda, MD) as follows. Thyroid gland size was measured as thyroid area on the photograph. Thyroid follicle size was measured using histological images at 50X magnification. The area of the thyroid gland was trimmed, converted to grayscale, and underwent black and white inversion. Particle analyses were performed on the processed images. Values greater than 500 μm^2^ were used. For each mouse, we evaluated the right and left lobes and calculated the average value. Follicular epithelial cell height was measured on histological images at 200X magnification. Heights at the upper, bottom, right, and left sides of follicles were measured. The average height from 5 follicles was calculated.

### Thyroid hormone measurement

Serum levels of fT3, fT4, and human TSH were measured using a FT3, FT4, TSH AccuLite VAST CLIA kit (Monobind Inc., Lake Forest, CA). We diluted serum samples 25-fold with phosphate buffered saline when we measured human TSH levels. Luminescence was measured using a 2030 ARVO X3 multilabel reader (PerkinElmer, Waltham, MA) or multilevel reader Spark (Tecan, Zurich, Switzerland).

### RNA extraction, RT-PCR, and quantitative RT-PCR

Total RNA was extracted using a Nucleospin RNA Plus kit (Macherey-Nagel) according to the manufacturer’s instructions. All extracted RNA was reverse transcribed using ReverTra Ace (TOYOBO Life Science, Osaka, Japan). Reverse transcription (RT)-PCR was performed using the Emerald Amp MAX PCR master mix (Takara Bio, Shiga, Japan). Quantitative RT-PCR was performed using THUNDERBIRD SYBR qPCR MIX (TOYOBO Life Science) with the StepOnePlus Real-time PCR System (Thermo Fisher Scientific), as in our previous studies (26, 27, 66, 67). Results were normalized using *Ppia* and *Hprt* as reference genes; relative mRNA expression of target genes was evaluated using the comparative threshold cycle method. Supplementary Table 4 lists the primers used.

### RNA-seq and bioinformatics analysis

The TruSeq Stranded mRNA Sample Prep Kit (Illumina, Inc., San Diego, CA) was used to construct mRNA paired-end libraries. Paired-end sequencing of 100 bp was performed on an NovaSeq 6000 system (Illumina, Inc.). Quality control metrics of raw sequencing reads were performed with FastQC version 0.11.7; low-quality reads were removed with Trimmomatic version 0.38. Next, reads were mapped to the mm10 reference sequence with HISAT2 version 2.1.0 and assembled into transcripts with StringTie version 2.1.3b. The expression profile was calculated as transcripts per kilobase million. For DEG analyses, fold change values were calculated and DEGs with an adjusted p[<[0.05 and fold changes[≤[−2 or[≥ 2 were determined using DESeq2. Gene-set enrichment analyses were performed based on GO with gProfiler and the KEGG database. These experiments and analyses were conducted by Macrogen Japan (Tokyo, Japan).

### Western blotting

We mechanically homogenized a lobe of the thyroid gland in 100 μL radioimmunoprecipitation buffer (Nacalai Tesque, Kyoto, Japan) and left the homogenate on ice for 30 minutes. Supernatants were centrifuged at 10,000 *g* for 10 minutes at 4°C and collected in microtubes. We evaluated protein concentrations using the Bradford method with Protein Assay CBB Solution (Nacalai Tesque); bovine serum albumin (Bio-Rad, Hercules, CA) was used as the standard. We performed electrophoresis with 20 μg/lane of each lysate in Bolt 4–12% Bis-Tris Plus Gels (Thermo Fisher Scientific). We transferred the protein samples onto polyvinylidene difluoride membranes with the iBlot2 Dry Blotting System (Thermo Fisher Scientific). The membranes were blocked with Bullet Blocking One (Nacalai Tesque). We incubated with primary antibodies overnight at 4°C followed by secondary antibodies for 2 hours at room temperature. Bands were detected using a chemiluminescent method with Chemi-Lumi One Super (Nacalai Tesque) in ImageQuant LAS 4000 (GE Healthcare, Chicago, IL). The primary antibody was a rabbit polyclonal anti-SLC26A4 antibody (PA5-115911; Thermo Fisher Scientific). A rabbit polyclonal anti-β-actin antibody (4967; Cell Signaling Technology, Danvers, MA) was used as the endogenous control. The secondary antibody was a horseradish-peroxidase-conjugated goat polyclonal anti-rabbit IgG antibody (4050-05; Southern Biotech, Cambridge, United Kingdom).

### Statistical analysis

All results are expressed as means ± standard error of the mean. Student’s *t*-test or one-way analysis of variance followed by the Dunnett test or the Tukey-Kramer test was used to analyze data on mouse phenotypes. JMP Pro version 17.0.0 (SAS Institute Inc., Cary, NC) was used for statistical analysis. Statistical analyses of DEGs were performed with the Wald test using DESeq2. Fisher’s exact test was used for enrichment analyses. Statistical significance was defined as *p* < 0.05.

## Acknowledgements

This work was supported by JSPS KAKENHI grants (Number 19K18006, 20K17508, 22K16394, and 23K15410) and by grants from Uehara Memorial Foundation, Takeda Science Foundation, and Japan Foundation for Applied Enzymology.

