## Supplementary Figures for "Transcriptomic Landscape of Hyperthyroidism in Mice Overexpressing Thyroid Stimulating Hormone"

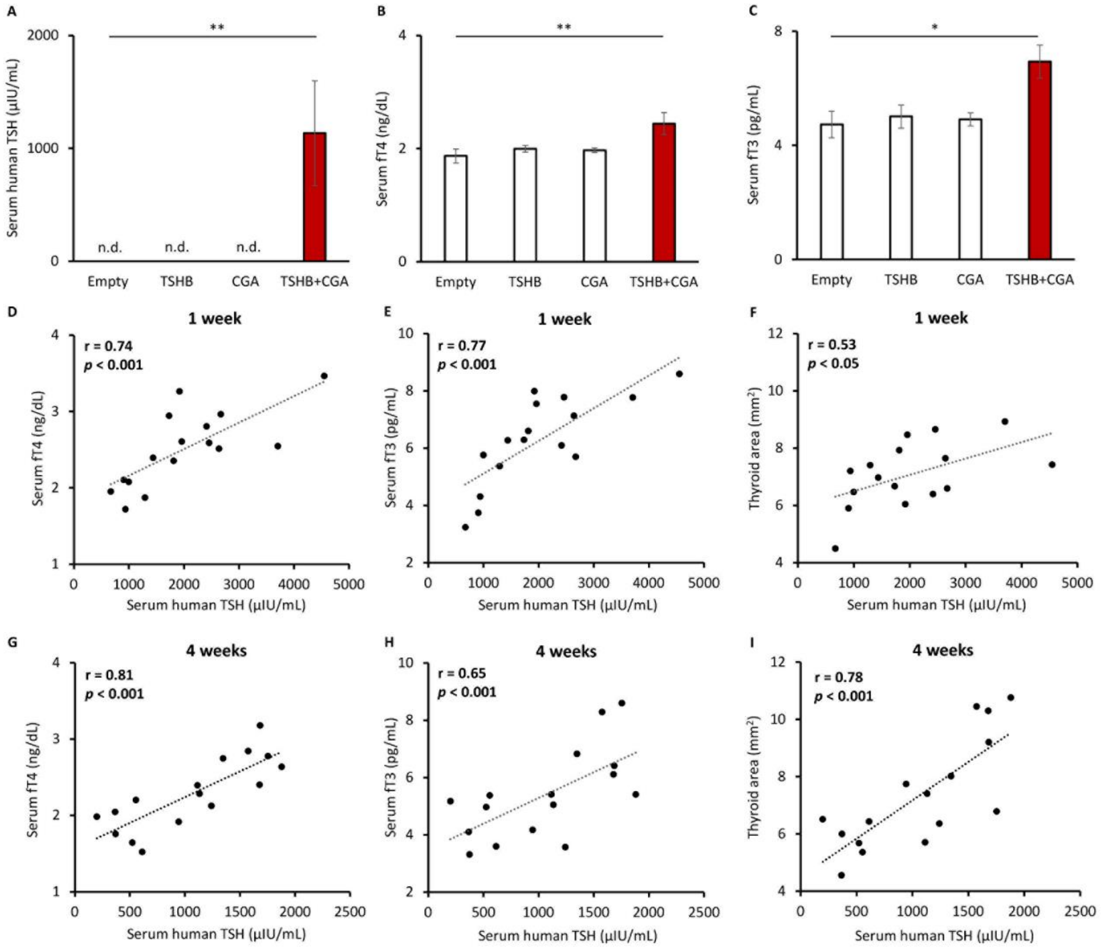


**Supplementary Figure 1.** (A–C) Serum levels of thyroid hormones of C57BL/6J mice injected with 25 μg of plasmid vectors; pLIVE-Empty (Empty), pLIVE-*TSHB* (TSHB), pLIVE-*CGA* (CGA), and pLIVE-*TSHB* and pLIVE-*CGA* (TSHB+CGA). n.d., not detected; TSH, thyroid stimulating hormone; fT4, free thyroxine; fT3, free triiodothyronine. Empty, TSHB, CGA, $n$ = 4 each, and TSHB+CGA, $n$ = 3. Data are represented as means ± standard error of the mean (SEM). One-way analysis of variance (ANOVA) followed by the Dunnett test with comparison to the Empty group. *$p < 0.05$ and **$p < 0.01$. (D–I) Correlation between serum human TSH levels and phenotypes of mice at 1 week (D–F, related to Figure 1) and at 4 weeks after hydrodynamic gene delivery (G–I, related to Figure 2). We plotted results of 8 mice of the 5 μg group and 8 mice of the 25 μg group together. Thyroid gland size was measured as thyroid area on photographs. Correlation was analyzed by calculating Pearson correlation coefficient.

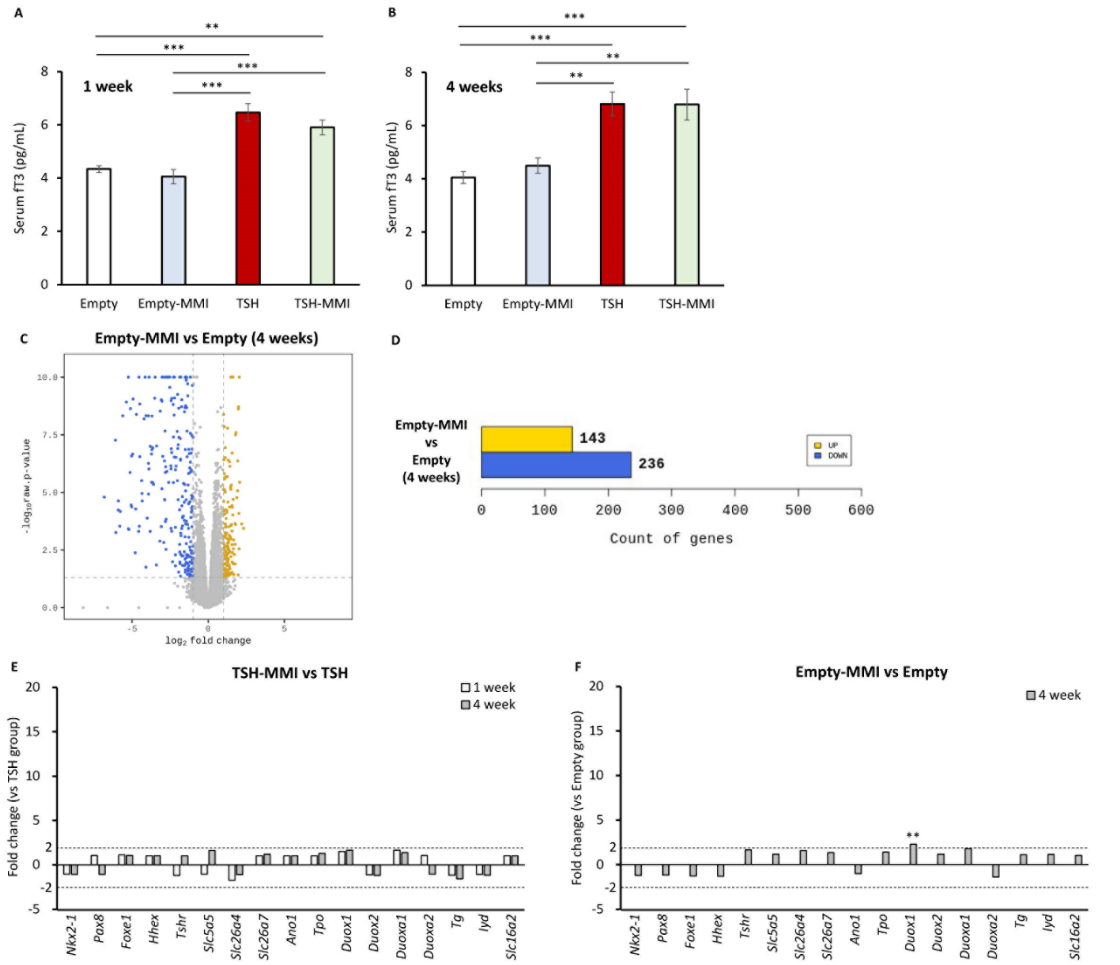


**Supplementary Figure 2.** (A, B) Serum fT3 levels of each group at 1 week (A, related to Figure 3) and at 4 weeks after hydrodynamic gene delivery (B, related to Figure 4). Data are represented as means ± SEM. Statistical analyses were performed using ANOVA followed by the Tukey-Kramer test. (C) Volcano plot of differentially expressed genes (DEGs) comparing the Empty-MMI and Empty group at 4 weeks (related to Figure 4), and (D) numbers of DEGs. (E, F) Summary of RNA sequencing (RNA-seq) regarding thyroidal genes. See Supplementary Tables 1 and 2 for the source data. Data are represented as fold changes in transcript per kilobase million (TPM) of genes. The Wald test was used to compare TPM of genes from each cohort. ***p* < 0.01 and ****p* < 0.001.

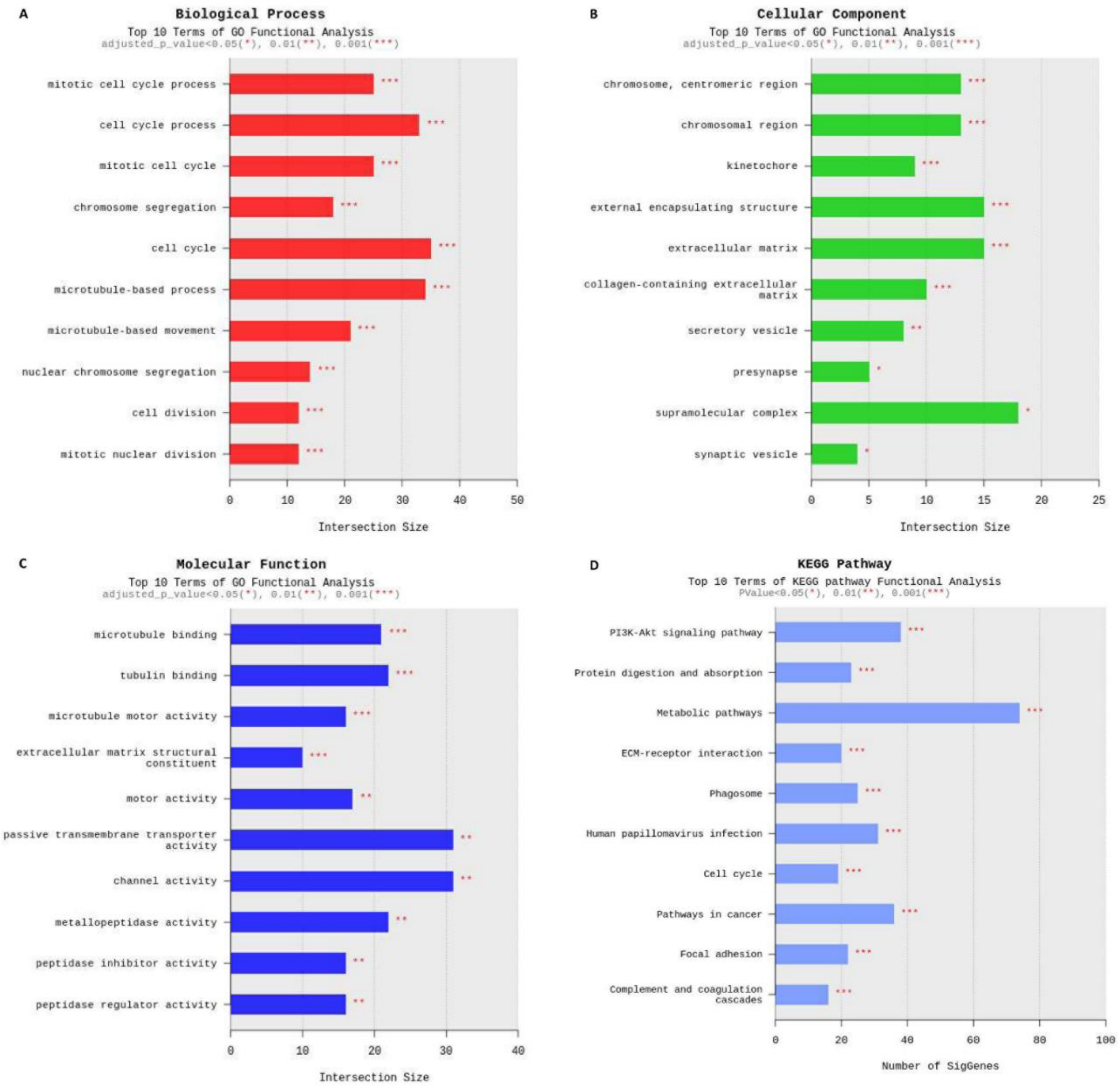


**Supplementary Figure 3.** Top 10 terms in enrichment analyses of the TSH group compared with the Empty group at 1 week after hydrodynamic gene delivery. (A–C) Gene Ontology (GO) terms. (D) Kyoto Encyclopedia of Genes and Genomes (KEGG) terms. Statistical analyses were performed using Fisher's exact test. *$p < 0.05$, **$p < 0.01$, and ***$p < 0.001$.

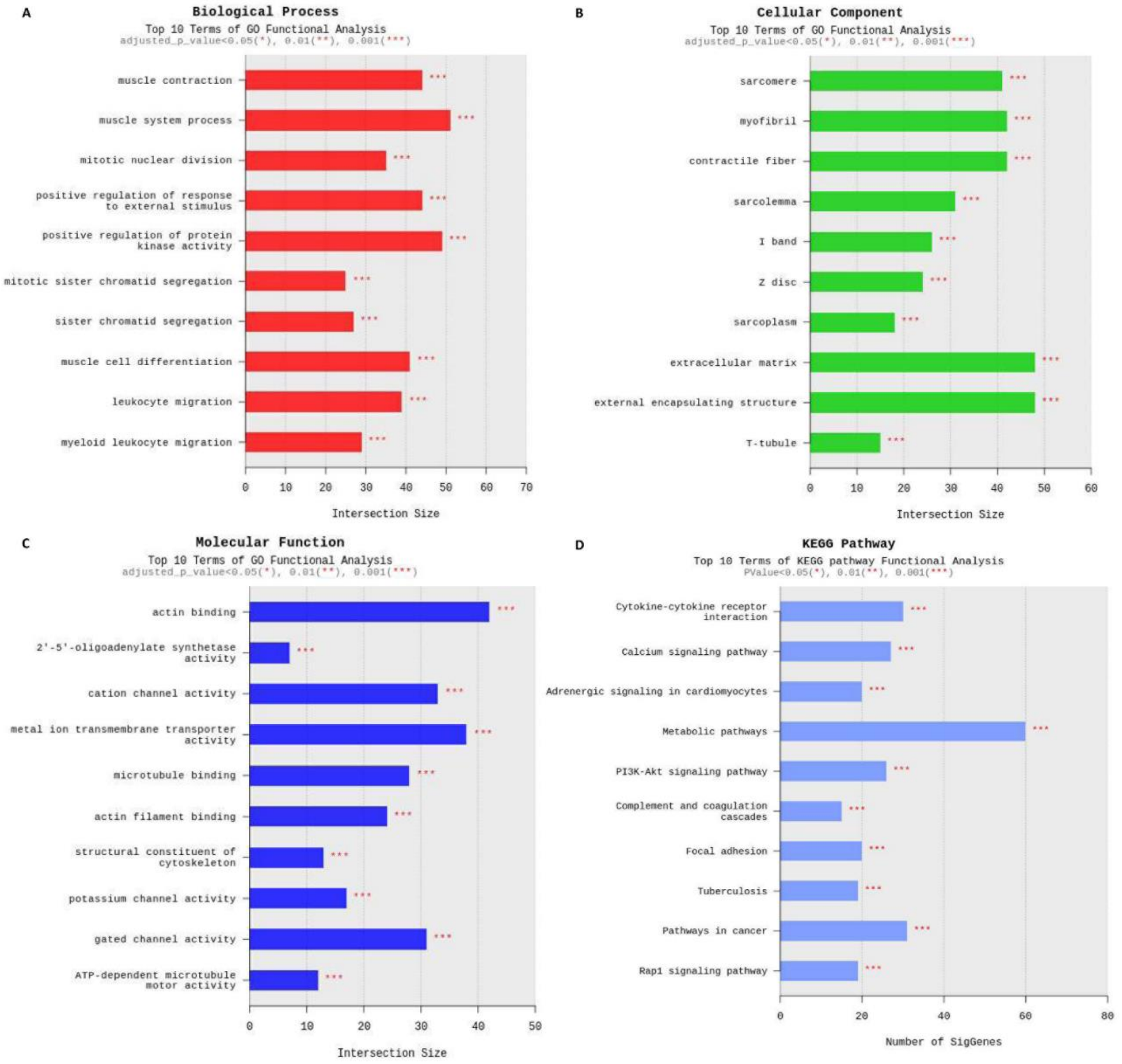


**Supplementary Figure 4.** Top 10 terms in enrichment analyses of the TSH group compared with the Empty group at 4 weeks after hydrodynamic gene delivery. (A–C) GO terms. (D) KEGG terms. Statistical analyses were performed using Fisher's exact test. ***$p < 0.001$.

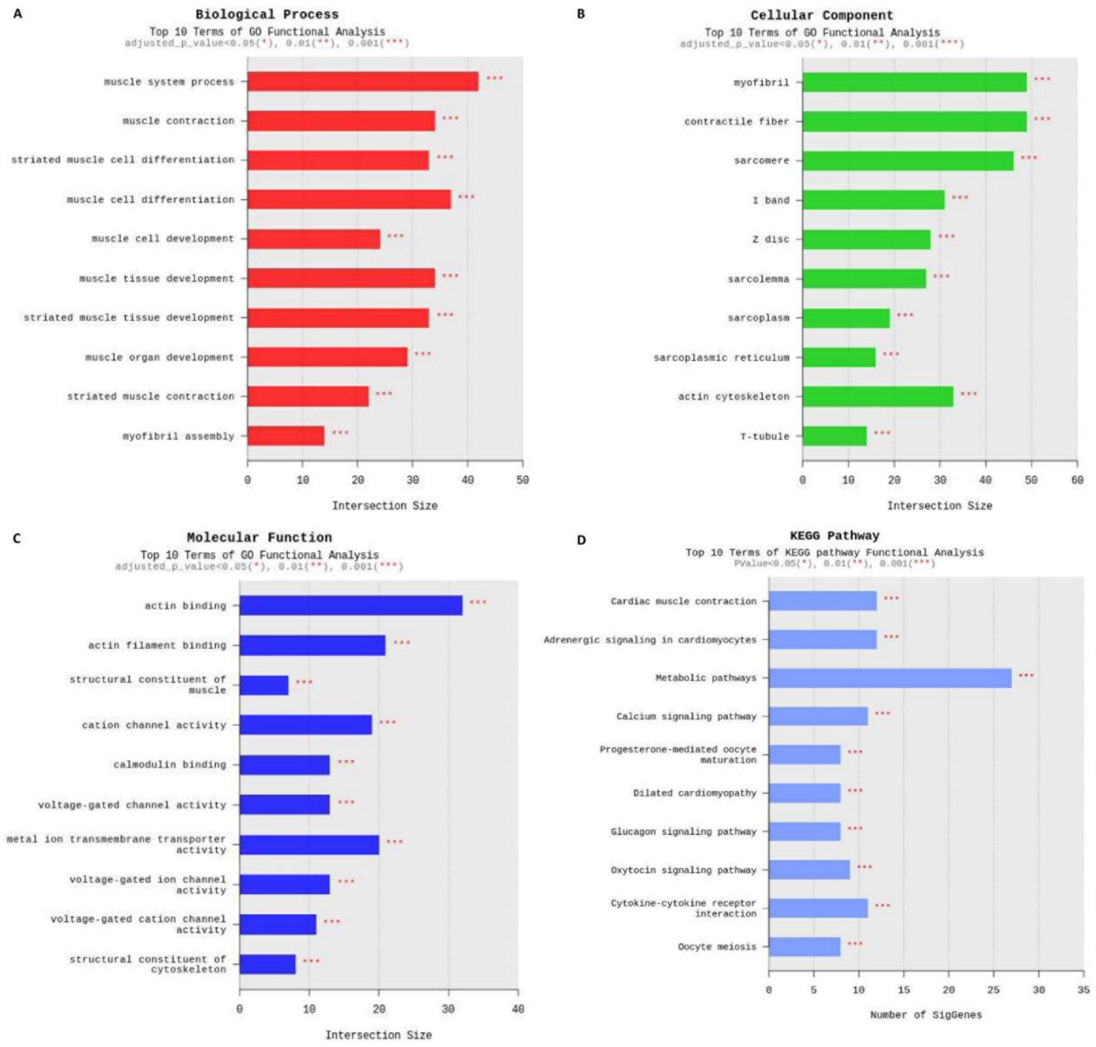


**Supplementary Figure 5.** Top 10 terms in enrichment analyses of the Empty-MMI group compared with the Empty group at 4 weeks after hydrodynamic gene delivery. (A–C) GO terms. (D) KEGG terms. Statistical analyses were performed using Fisher's exact test. ***$p < 0.001$.

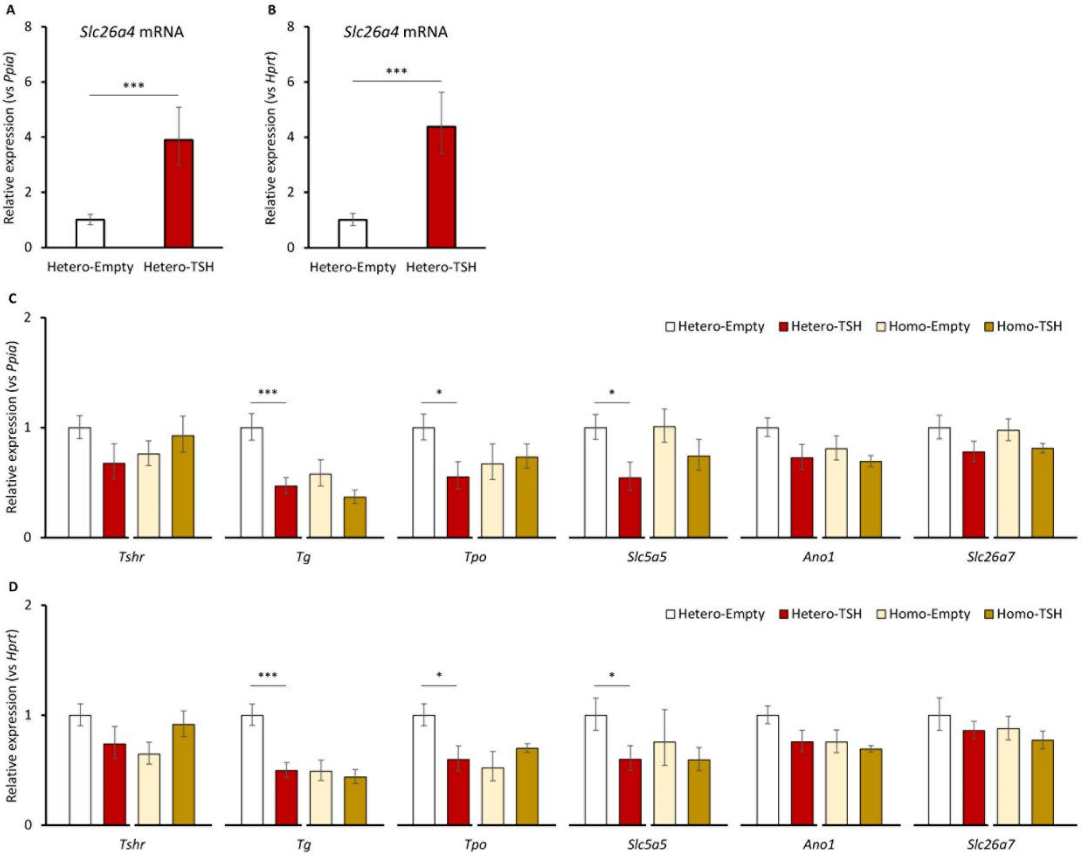


**Supplementary Figure 6.** Analyses of the thyroid glands of *Slc26a4* knockout mice at 1 week after hydrodynamic gene delivery (related to Figure 6). (A, B) *Slc26a4* mRNA level of the thyroid gland determined by quantitative reverse transcription-polymerase chain reaction (RT-PCR). Results are normalized using *Ppia* (A) and *Hprt* (B) as internal controls. (C, D) Expressions of genes related to thyroid hormone secretion and iodine transport determined by quantitative RT-PCR. Results are normalized using *Ppia* (C) and *Hprt* (D) as internal controls. Hetero-Empty, Hetero-TSH, $n = 9$ each; Homo-Empty, Homo-TSH, $n = 5$ each. Data are represented as means ± SEM. Statistical analyses were performed using Student's t-test to compare the Empty and TSH groups for each knockout mouse. $^{*}p < 0.05$ and $^{***}p < 0.001$.

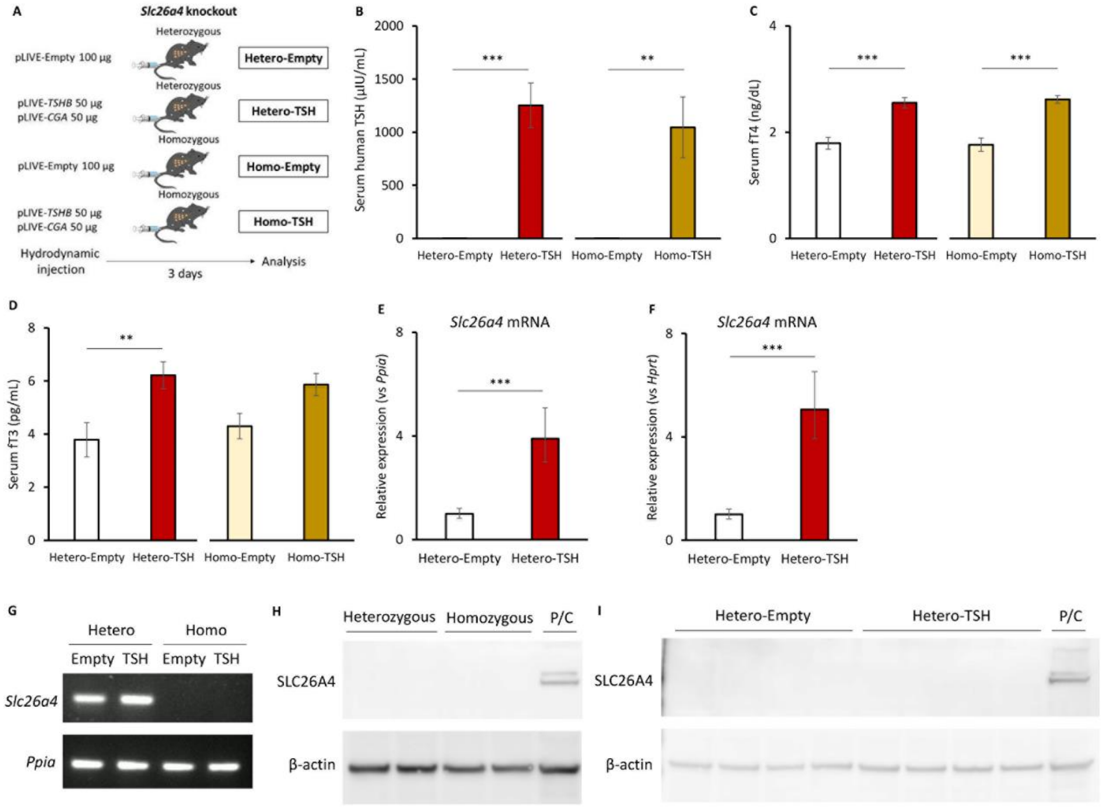


**Supplementary Figure 7.** Thyroid function and SLC26A4 expressions of *Slc26a4* knockout mice at 3 days after hydrodynamic gene delivery. (A) Schema for the experimental design. (B–D) Serum levels of thyroid hormones. (E, F) *Slc26a4* mRNA level of the thyroid gland determined by quantitative RT-PCR. Results are normalized using *Ppia* (E) and *Hprt* (F) as internal controls. (G) *Slc26a4* mRNA expressions determined by RT-PCR. (H, I) Western blot analyses of SLC26A4 expressions in the thyroid glands. P/C means a positive control of cerebral lysate of heterozygous *Slc26a4* knockout mice. SLC26A4 was detected as a lower band at a molecular weight of 90,000. Hetero-Empty, Hetero-TSH, $n = 4$ each; Homo-Empty, Homo-TSH, $n = 3$ each. Data are represented as means ± SEM Statistical analyses were performed using Student's t-test to compare the Empty and TSH groups for each knockout mouse. *$p < 0.05$ and ***$p < 0.001$.

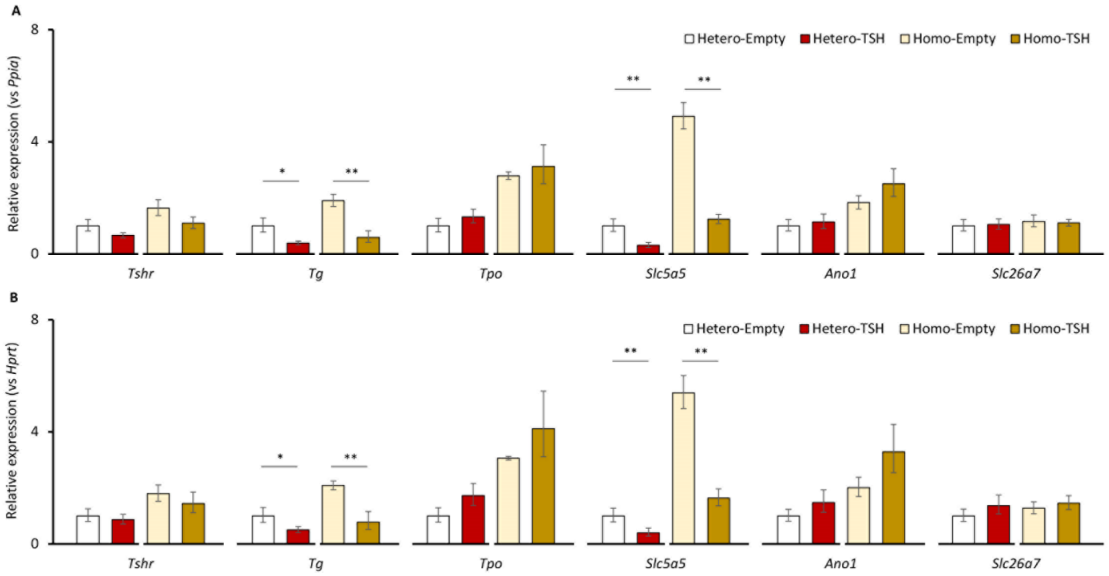


**Supplementary Figure 8.** Analyses of the thyroid glands of *Slc26a4* knockout mice at 3 days after hydrodynamic gene delivery. (A, B) Expressions of genes related to thyroid hormone secretion and iodine transport determined by quantitative RT-PCR. Results are normalized using *Ppia* (A) and *Hprt* (B) as internal controls. Hetero-Empty, Hetero-TSH, $n = 4$ each; Homo-Empty, Homo-TSH, $n = 3$ each. Data are represented as means ± SEM. Statistical analyses were performed using Student's t-test to compare the Empty and TSH groups for each knockout mouse. *$p < 0.05$ and **$p < 0.01$.
