## Supplementary Table 4 for "Transcriptomic Landscape of Hyperthyroidism in Mice Overexpressing Thyroid Stimulating Hormone"

| **Supplementary Table 4. Primers used for plasmid construction, quantitative RT-PCR, and RT-PCR** | | |
| --- | --- | --- |
| **Genes (accession number)** | Fw: forward primer (5′–3′) | Rv: reverse primer (5′–3′) |
| **Plasmid construction** | | |
| Human *TSHB* (NM_000549) | AATAGCTAGCGCCACCATGACTGCTCTCTTTCTG | CTGGTAGGATTTTCTGTCTAAGGATCCTATT |
| Human *CGA* (NM_000735) | AATAGCTAGCGCCACCATGGATTACTACAGAAAAT | CTTGTTATTATCACAAATCTTAAGGATCCTATT |
| **Quantitative RT-PCR** | | |
| *Slc26a4* (NM_011867) | GCTCGCATTCGGGACTGTAA | CAGCAAACCTGCTTTGGCAT |
| *Tshr* (NM_011648) | TCCCTGAAAACGCATTCCA | GCATCCAGCTTTGTTCCATTG |
| *Tg* (NM_009375) | TGTCCCACCAAGTGTGAAAA | CCAAGGAAAGCTTGTTCAGC |
| *Tpo* (NM_009417) | CAAAGGCTGGAACCCTAATTTCT | AACTTGAATGAGGTGCCTTGTCA |
| *Slc5a5* (NM_053248) | GGGATGCACCAATGCCTCTG | GTAGCTGATGAGAGCACCACA |
| *Slc26a7* (NM_145947) | CCCCACCGAGAAGACATTAAGC | TGAACTGCCAACATTATCCCAG |
| *Ano1* (NM_178642) | GAGGCCAGTAGCCATCAGAG | CGTGAAGGAGATCACAAAGGC |
| *Ppia* (NM_008907) | CGCGTCTCCTTCGAGCTGTTTG | TGTAAAGTCACCACCCTGGCACAT |
| *Hprt* (NM_013556) | GGACCTCTCGAAGTGTTGGATAC | GCTCATCTTAGGCTTTGTATTTGGCT |
| **RT-PCR** | | |
| *Slc26a4* (NM_011867) | ATTGCTACTGCCATTTCCTATG | AGGATGCAGCCAACATGTCCG |
| *Ppia* (NM_008907) | CGCGTCTCCTTCGAGCTGTTTG | TGTAAAGTCACCACCCTGGCACAT |

RT-PCR, reverse transcription-polymerase chain reaction.
